# Accurate evaluation of live-virus microneutralization for SARS-CoV-2 variant JN.1

**DOI:** 10.1101/2024.04.17.589886

**Authors:** Giulia Dowgier, Agnieszka Hobbs, David Greenwood, Marianne Shawe-Taylor, Phoebe Stevenson-Leggett, James Bazire, Rebecca Penn, Ruth Harvey, Crick COVID serology pipeline, Legacy Investigators, Vincenzo Libri, George Kassiotis, Steve Gamblin, Nicola S Lewis, Bryan Williams, Charles Swanton, Sonia Gandhi, David LV Bauer, Edward J Carr, Emma C Wall, Mary Y Wu

## Abstract

Emerging SARS-CoV-2 variants require rapid assessments of pathogenicity and evasion of existing immunity to inform policy. A crucial component of these assessments is accurate estimation of serum neutralising antibody titres using cultured live virus isolates. Here, we report our updated culture methods for Omicron sub-variant JN.1 using Caco-2 cells and the subsequent evaluation of neutralising antibody titres (nAbTs) in recipients of BNT162b2-XBB.1.5 monovalent and the Ancestral/BA.5 containing bivalent vaccines. We compared culture of JN.1 in either Vero V1 cells or Caco-2 cells, finding culture in Vero V1 either resulted in low-titre stocks or induced crucial mutations at the Spike furin cleavage site. Using the sequence-clean culture stocks generated in Caco-2 cells, we assessed serum samples from 71 healthy adults eligible for a COVID-19 vaccination given as a 5^th^ dose booster: all participants had detectable nAbs against JN.1 prior to vaccination, with baseline/pre-existing nAbTs between both vaccine groups comparable (p = 0.240). However, nAbTs against JN.1 post-vaccination were 2.6-fold higher for recipients of the monovalent XBB1.5 vaccine than the BA.4/5 bivalent vaccine (p<0.001). Regular re-appraisal of methods involved in the evaluation of new variants is required to ensure robust data are used to underpin crucial severity assessments as variants arise and vaccine strain selection decisions.

## Main text

Since the emergence of omicron BA.1 in 2021, SARS-CoV-2 omicron lineage sub-variants continue to dominate the global COVID-19 landscape. Emerging mutations in the Spike protein confer enhanced replication and transmissibility of these sub-variants in the face of increasing population ‘hybrid’ immunity^1^. In response, COVID-19 mRNA vaccines have been updated three times, but the response to new variants, including vaccine strain selection decisions and manufacturing capabilities mean updated vaccines are delivered at least 6-12 months behind each variant emergence^2^. Further, a lack of prospective modelling to determine pre-existing population immunity/correlates of protection means that estimates of severe COVID risk in both clinically vulnerable and the wider population requires rapid *in vitro* characterisation of new variants by neutralisation of sera. Such data remains essential to evaluate immune escape, relative boosting efficacy of updated vaccines and inform strain-selection decisions.

The Crick’s COVID Surveillance Unit (CSU) hosts a state-of-the-art high throughput live-virus microneutralisation assay developed in 2020, enabling rapid, accurate characterisation of thousands of serum samples against multiple live variants in near real-time. We have provided data to help calibrate WHO standards for neutralisation^3,4^, derived correlates of protection for alpha and delta variants^5,6^ and generated data on vaccine-induced neutralisation for healthy adults and immunocompromised individuals - including patients with cancer and kidney failure - that have informed policy^7-16^. Our assay uses live viruses isolated from research participants with COVID-19 infection through the UCLH-Crick Legacy study (NCT04750356) in the United Kingdom, and from collaborators through our participation in wider networks including the Genotype-to-Phenotype 2 UK consortium. Virus stocks undergo stringent quality control (QC) checks to ensure that they are free of culture-induced mutations and are of sufficient titres for the microneutralisation assay (Fig 1A).

**Figure 1:**
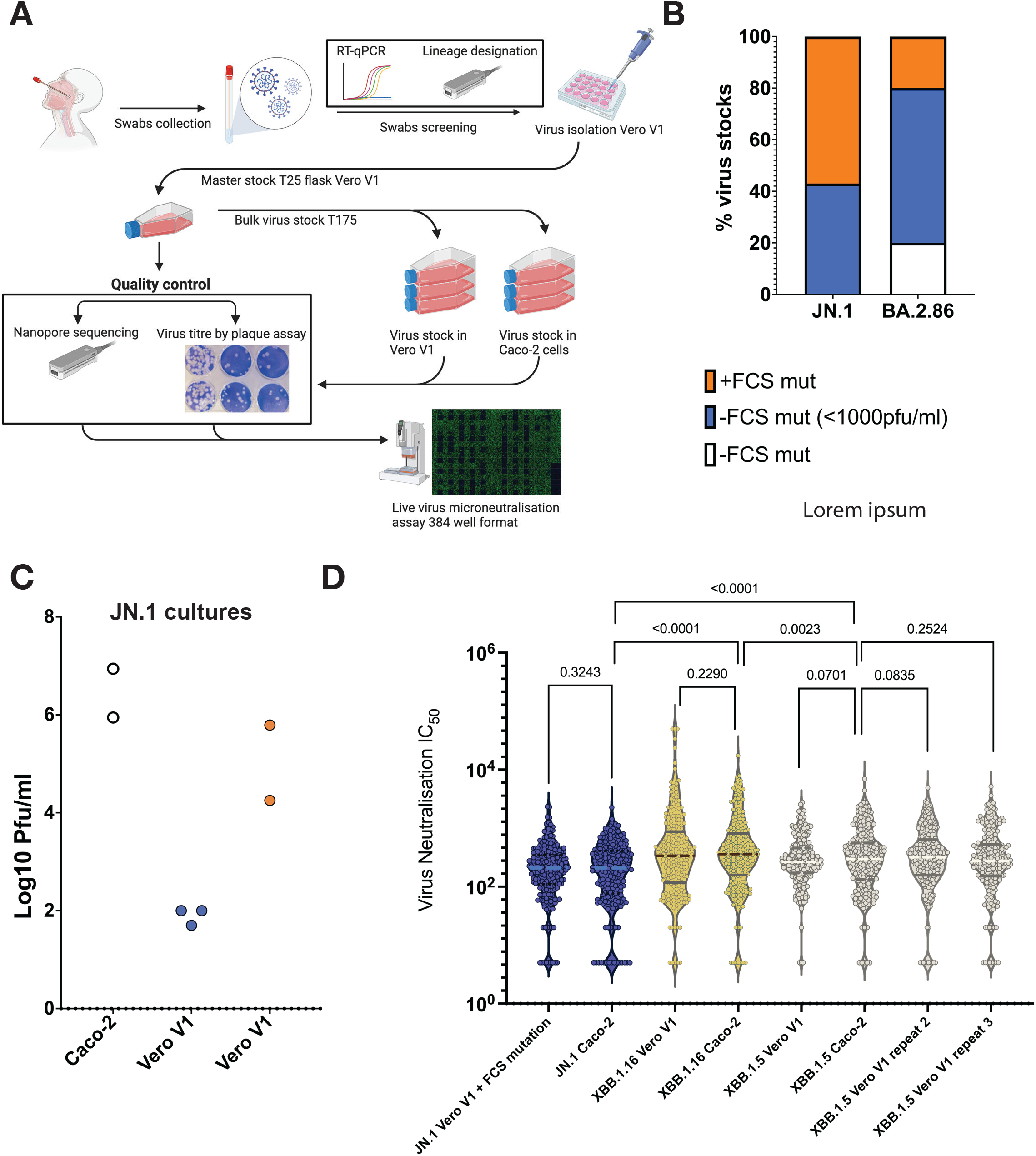
Furin cleavage site mutations in SARS-CoV-2 Spike are induced by culturing in VeroV1 cells and not in Caco-2 cells and do not affect neutralisation assay results. (**A)** Schematic of the steps involved in The Crick’s COVID Surveillence Unit (CSU) live virus isolation and culture for use in microneutralisation assays. (**B)** Proportion of virus stocks cultured in Vero V1 cells containing an FCS mutation (orange) is higher in JN.1 than BA.2.86. (**C**) JN.1 cultured in Caco2 cells reach high titres without acquiring FCS mutations (white) compared to culture in Vero V1 cells where high titres are only achieved in isolates acquiring FCS mutations (+ mutation orange, n=2, -mutation blue, n=3), y axis = Log10 of plaque forming units/ml (Pfu/ml). (**D**) nAbTs for a reference panel of 214 serum samples were measured against 6 different virus stocks comprising 3 different variants grown in either Vero V1 or Caco-2 cells. JN.1 grown in Vero V1 cells acquired a mutation in the FCS. Neither cell line nor FCS mutations affected nAbTs for the same variant (P > 0.07 for all comparisons), whereas comparison of JN.1 to either XBB.1.5 or XBB.1.16 results in statistically different titres for the same samples (P < 0.0001) for both comparisons). XBB.1.5 and XBB.1.16 differ by 1 mutation in Spike and this is reflected in similar nAbTs (p = 0.0023). The XBB.1.5 stock grown in Vero V1 cells was used for 3 total biological repeats on different days to assess reproducibility of the assay. P values shown calculated using two-tailed Wilcoxon tests.

Omicron BA.2.86, and its daughter variant JN.1 caused a substantial global wave of COVID-19 across late 2023/early 2024. We recently reported on the divergent neutralisation response in healthy adults to two different COVID-19 mRNA vaccines, given as a 5^th^ dose in the UK’s autumn 2023 booster campaign^7^. Following isolation of JN.1, we prepared to extend our original comparison between the two different vaccines using this new variant.

However, a JN.1 virus stock passing QC required a significant adaptation to our existing culture protocols. Here, we report our updated culture methods for JN.1 using Caco-2 cells and the subsequent evaluation of neutralising antibody titres (nAbTs) in recipients of BNT162b2-XBB.1.5 monovalent and the Ancestral/BA.5 containing bivalent vaccines.

Viral culture for the Crick’s microneutralisation assay has previously been undertaken in Vero V1 cells^17,18^, which express the parainfluenza virus 5 (PIV5) V protein^19^. We previously found that viruses cultured in these Vero V1 cells were less likely to acquire cell culture adaptive mutations as compared to “conventional” Vero cell lines, in which we and others^20-24^ have found that SARS-CoV-2 was more prone to accumulating mutations at the furin cleavage site (FCS) in the Spike protein. (Fig 1B). However, we found that JN.1 isolates were highly unstable, even in Vero V1 cells: JN.1 sequence-clean stocks tend to have very low or unusable titres relative to those with FCS mutations (Fig 1B and C) suggesting selective pressure in Vero V1 cells for the accumulation of these mutations.

To obtain a clean stock of JN.1 for microneutralisation assays, we tried passaging virus stocks in Caco-2 cells and were able to obtain high viral titres with clean sequences (Fig 1C). We tested a reference panel of 214 serum samples from Legacy participants spanning vaccine doses 1-4 against three recent variants (XBB.1.5, XBB.1.6, and JN.1), each with stocks grown in both Vero V1 and Caco-2 cells — with JN.1 in Vero V1 cells having acquired a FCS mutation (S:R685H). There is no significant difference in nAbTs between virus stocks passaged in Vero V1 cells and Caco-2 cells, nor with and without FCS mutations, for any of the intra-variant comparisons (Fig 1D, p > 0.07). Whereas, nAbTs for comparisons between different variants are significantly different as expected (Fig 1D, p<0.0023).

We then analysed 140 serum samples from 71 Legacy participants who received a 5^th^ dose of the SARS-CoV-2 vaccine (Table 1). We found that when stratified by the type of vaccine administered, the older bivalent ancestral/BA.5 vaccine does not provide as large of a boost in nAbTs against JN.1 as the newer monovalent XBB.1.5 vaccine (Fig 2A). We previously found that there was no difference in the nAbT boosting effect of either vaccine against either Ancestral virus or BA.2.86, despite divergent responses towards other variants^7^. Most participants had detectable nAbs against JN.1 prior to vaccination, with baseline/pre-existing nAbTs between both vaccine groups comparable (Fig 2B, p = 0.240). However, nAbTs against JN.1 post-vaccination were 2.6-fold higher for the monovalent XBB1.5 vaccine than the BA.4/5 bivalent vaccine (Fig 2B, p < 0.001). This is surprising based on our previous findings against BA.2.86, since JN.1 has just 1 additional mutation relative to the BA.2.86 Spike (S:L455S) (Fig 2, and ^7^).

**Table 1.**
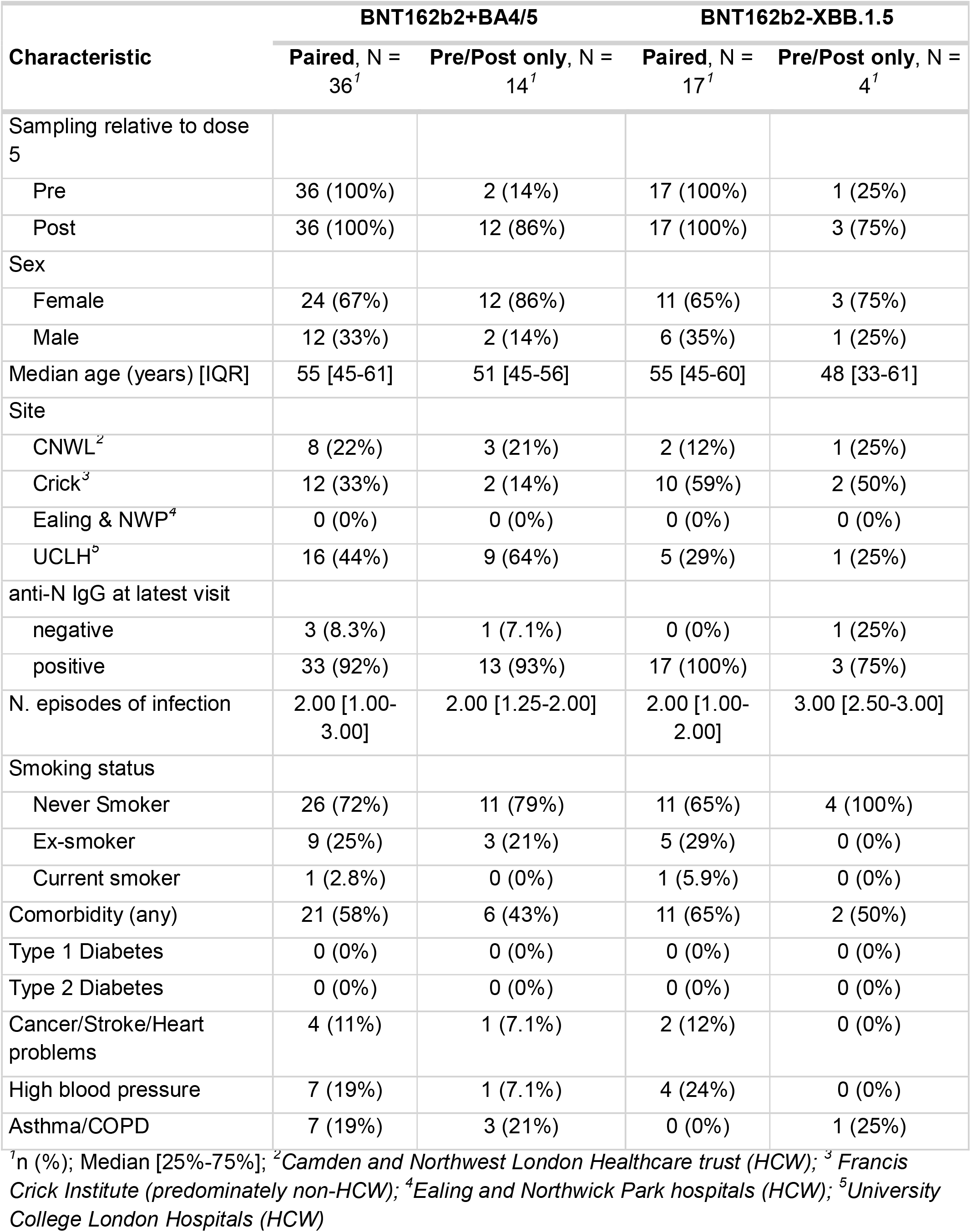
Description of cohort characteristics grouped by fifth dose vaccine type and those sampled pre- and post-vaccination.

| Characteristic | BNT162b2+BA4/5 |  | BNT162b2-XBB.1.5 |  |
| --- | --- | --- | --- | --- |
|  | Paired, N =<br>36<sup>1</sup> | Pre/Post only, N =<br>14<sup>1</sup> | Paired, N =<br>17<sup>1</sup> | Pre/Post only, N =<br>4<sup>1</sup> |
| Sampling relative to dose 5 |  |  |  |  |
| Pre | 36 (100%) | 2 (14%) | 17 (100%) | 1 (25%) |
| Post | 36 (100%) | 12 (86%) | 17 (100%) | 3 (75%) |
| Sex |  |  |  |  |
| Female | 24 (67%) | 12 (86%) | 11 (65%) | 3 (75%) |
| Male | 12 (33%) | 2 (14%) | 6 (35%) | 1 (25%) |
| Median age (years) [IQR] | 55 [45-61] | 51 [45-56] | 55 [45-60] | 48 [33-61] |
| Site |  |  |  |  |
| CNWL<sup>2</sup> | 8 (22%) | 3 (21%) | 2 (12%) | 1 (25%) |
| Crick<sup>3</sup> | 12 (33%) | 2 (14%) | 10 (59%) | 2 (50%) |
| Ealing & NWP<sup>4</sup> | 0 (0%) | 0 (0%) | 0 (0%) | 0 (0%) |
| UCLH<sup>5</sup> | 16 (44%) | 9 (64%) | 5 (29%) | 1 (25%) |
| anti-N IgG at latest visit |  |  |  |  |
| negative | 3 (8.3%) | 1 (7.1%) | 0 (0%) | 1 (25%) |
| positive | 33 (92%) | 13 (93%) | 17 (100%) | 3 (75%) |
| N. episodes of infection | 2.00 [1.00-3.00] | 2.00 [1.25-2.00] | 2.00 [1.00-2.00] | 3.00 [2.50-3.00] |
| Smoking status |  |  |  |  |
| Never Smoker | 26 (72%) | 11 (79%) | 11 (65%) | 4 (100%) |
| Ex-smoker | 9 (25%) | 3 (21%) | 5 (29%) | 0 (0%) |
| Current smoker | 1 (2.8%) | 0 (0%) | 1 (5.9%) | 0 (0%) |
| Comorbidity (any) | 21 (58%) | 6 (43%) | 11 (65%) | 2 (50%) |
| Type 1 Diabetes | 0 (0%) | 0 (0%) | 0 (0%) | 0 (0%) |
| Type 2 Diabetes | 0 (0%) | 0 (0%) | 0 (0%) | 0 (0%) |
| Cancer/Stroke/Heart problems | 4 (11%) | 1 (7.1%) | 2 (12%) | 0 (0%) |
| High blood pressure | 7 (19%) | 1 (7.1%) | 4 (24%) | 0 (0%) |
| Asthma/COPD | 7 (19%) | 3 (21%) | 0 (0%) | 1 (25%) |

<sup>1</sup>n (%); Median [25%-75%]; <sup>2</sup>Camden and Northwest London Healthcare trust (HCW); <sup>3</sup>Francis Crick Institute (predominately non-HCW); <sup>4</sup>Ealing and Northwick Park hospitals (HCW); <sup>5</sup>University College London Hospitals (HCW)

**Figure 2:**
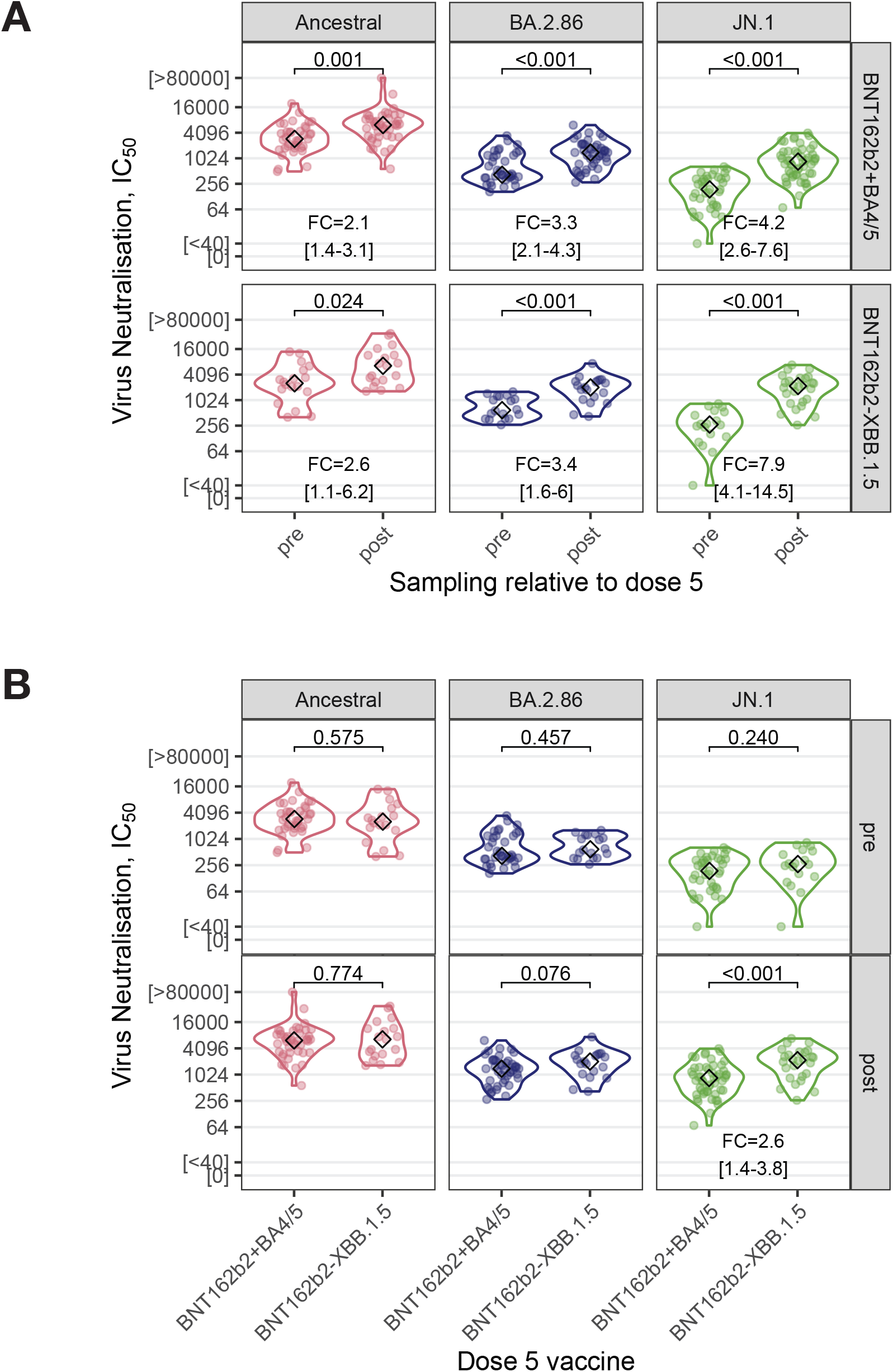
Recipients of BNT162b2-XBB.1.5 monovalent vaccine given as a 5th dose boosts serum neutralisation of JN.1 better than the Ancestral/BA.4/5 bivalent vaccine. (**A)** Distribution of live-virus nAbTs against SARS-CoV-2 Ancestral, BA.2.86, and JN.1 subvariants across the cohort are shown as the log2 of the IC50 for serum samples taken before or after a 5th dose vaccination with the bivalent BNT162b2 ancestral+BA.4/5 (top row) or BNT162b2-XBB.1.5 monovalent vaccine (bottom row). **(B)** nAbTs stratified for pre-vaccination (top row) and post-vaccination (bottom row) comparing the vaccines. p values shown are from unpaired, two-tailed Wilcoxon tests, or McNemar’s χ2 tests if the median of one group was more or less than the quantitative range of the assay (40–2560). FC = fold-change increase of nAbTs with 95% CIs in brackets. IC50 = 50% inhibitory concentration.

To verify that our JN.1 result is not an artifact from a cell line-specific adaptation, we repeated using sequence-clean JN.1 stocks grown in both Vero V1 (with low titres) and Caco-2 cells (Fig 3). While the observed range of nAbTs against JN.1 grown in Vero V1 cells is slightly compressed relative to JN.1 grown in Caco-2 cells (Fig S1A), the correlation is high (Fig S1B, C, and D), and differential boost by the monovalent and bivalent vaccines holds true (Fig 3A and B, FC between 2.6 and 3.2, p < 0.001). The low viral titre of the Vero V1 JN.1 stock meant a larger proportion of the harvested supernatant was used during infection, and potentially cell debris or incomplete viral particles not removed during clarification may be more enriched which could lead to the observed effects on nAbTs.

**Figure 3:**
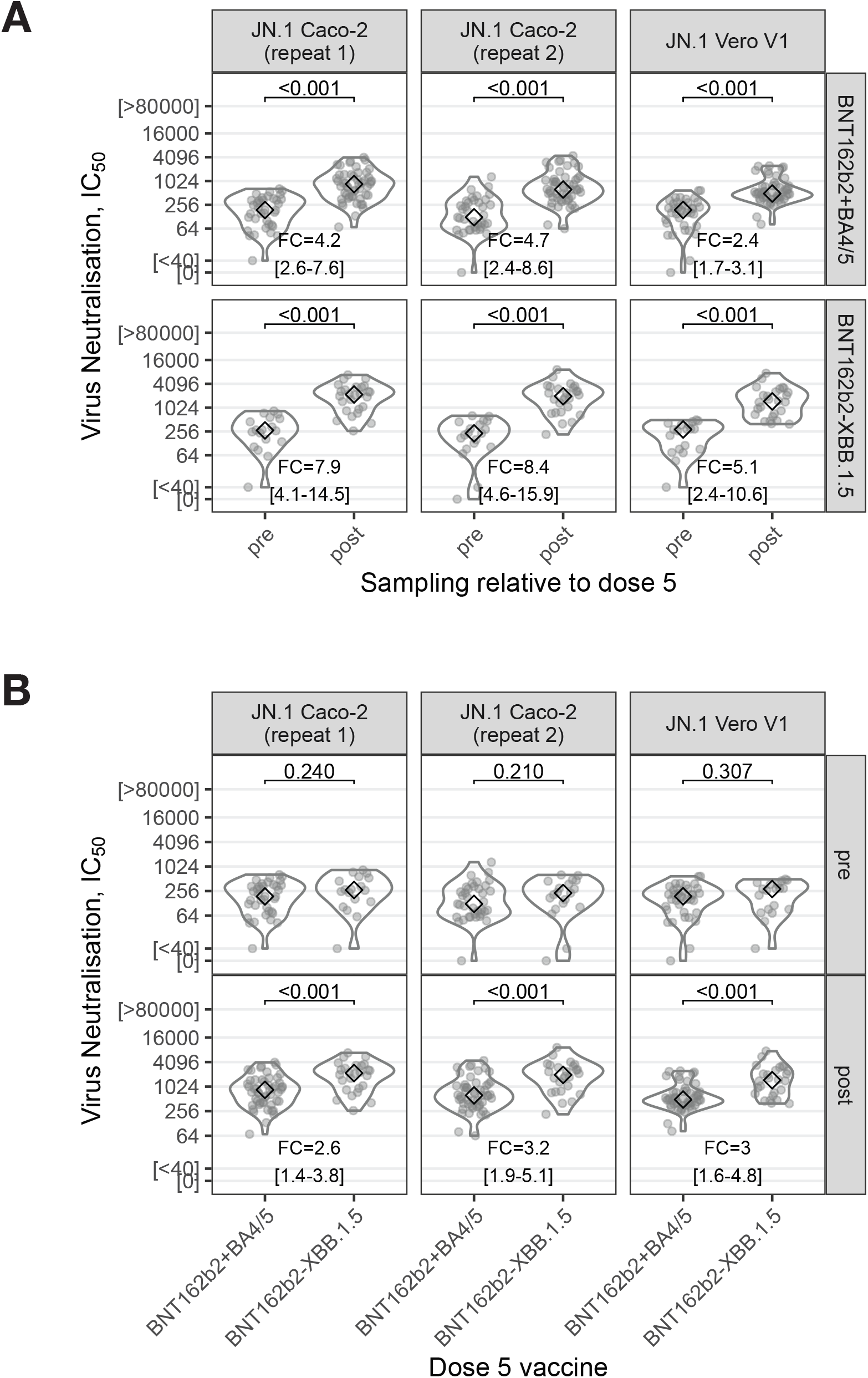
JN.1 cultured in Vero V1 and Caco-2 cells show the same differential boost of nAbTs by monovalent XBB.1.5 and bivalent Ancestral/BA.4/5 vaccines. (**A)** Distribution of live-virus microneutralisation titres against SARS-CoV-2 JN.1 cultured in Caco-2 cells in two separate experiments and Vero V1 cells are shown as the log2 of the IC50 for serum samples taken before or after a 5th dose of either the bivalent BNT162b2 ancestral+BA.4/5 (top row) or BNT162b2-XBB.1.5 monovalent vaccine (bottom row). **(B)** stratified by pre-vaccination (top row) and post-vaccination (bottom row) for comparisons between vaccines. p values shown are from unpaired, two-tailed Wilcoxon tests, or McNemar’s χ2 tests if the median of one group was more or less than the quantitative range of the assay (40–2560). FC = fold-change increase of nAbTs with 95% CIs in brackets. IC50 = 50% inhibitory concentration.

Our results highlight important considerations for the study of live SARS-CoV-2 virus. First, while we observe no difference in nAbTs between JN.1 stocks with and without FCS mutations, an intact FCS is critical for SARS-CoV-2 pathogenesis^22^. Therefore, detailed sequence verification of stocks (including examination of minor allele frequencies) will be especially critical for *in vivo* assessment of JN.1 pathogenicity in animal models. Second, culturing techniques need to be adapted to emerging variants specifically to avoid the use of low titre stocks, which may affect readouts even if the virus is sequence-validated.

Importantly, regular re-appraisal of methods involved in the evaluation of new variants is required to ensure robust data are used to underpin crucial severity assessments as variants arise and vaccine strain selection decisions.

## Main Table

## Methods

### Clinical cohort

The UCLH-Crick Legacy study (NCT04750356) is a prospective observational cohort, established in January 2021 to investigate longitudinal immunity to SARS-CoV-2. Extensive descriptions of the cohort can be found in our prior reports^7-10,14,15,25^. In the UK, from September 2023, healthcare workers, adults over 65 years, and those with either immunocompromised or caring responsibilities were offered a dose of COVID-19 vaccine as a 5^th^ dose booster. The majority of Legacy study participants eligible for this campaign received either a dose of bivalent COVID-19 vaccine containing mRNA encoding Ancestral and Omicron BA.4/5 Spikes (BNT162b2+BA4/5) or a monovalent COVID-19 vaccine containing mRNA encoding the XBB.1.5 Spike (BNT162b2-XBB.1.5) as a fifth dose. Participants were invited for paired pre- and post-vaccination visits approximately 1 week before and 3 weeks after the dose. If an individual was unable to attend pre-vaccination, their dose was not delayed. At each study visit, individuals performed a nasopharyngeal swab into virus transport medium (VTM; MWE Sigma-Virucult), gave details on any recent infection episodes, and had blood drawn for serum for live-virus microneutralisation assays and anti-N IgG detection.

Legacy participants were included in this study if they received their fifth dose of COVID-19 vaccine (BNT162b2+BA4/5 or BNT162b2-XBB.1.5) after August 1st 2023 and had a pre-boost sample taken more than 2 weeks after a previous dose and/or a post-boost sample within 4 weeks of a fifth dose^7^. We also analysed a subset of participants who contributed paired pre- and post-vaccination serum samples^7^.

### Virus variants and culture

All viral isolates were propagated in Vero V1 or Caco-2 cells. Briefly, 50-75% confluent monolayers of Vero V1 or Caco-2 cells were infected with the given SARS-CoV-2 variant at an MOI of approx. 0.001. Cells were washed once with PBS, then 5 ml virus inoculum made up in DMEM + 1% FCS was added to each T175 flask and incubated at room temperature for 1h. DMEM + 1% FCS was then added to each flask. Cells were incubated at 37° C, 5% CO2 for 3-4 days. To monitor viral growth during incubation, supernatant is sampled and RNA is extracted and purified (Qiagen Viral RNA mini kit) before quantification of viral genome number via RT-qPCR (TaqPath COVID-19 CE-IVD Kit, ThermoFisher). Final supernatant was harvested and clarified by centrifugation at 4000 rpm for 10 minutes in a benchtop centrifuge then aliquoted and frozen at −80°C.

Whole viral genomes were sequenced using a Nanopore MinION R10 flow cell following RT PCR expansion using the Midnight kit (EXP-MRT001) and barcoding using the Rapid Barcoding kit (SQK-RBK114.96) from purified RNA (Qiagen Viral RNA mini kit). For screening of Legacy participants infected with SARS-CoV-2 and further lineage designation, RNA was extracted and purified from nasopharyngeal swabs using the Qiagen Viral RNA mini kit following manufacturer’s instructions. RNA extracted from swabs underwent RT-qPCR analysis (TaqPath COVID-19 CE-IVD Kit, ThermoFisher) to confirm SARS-CoV-2 infection and assess the presence of S-gene target failure (SGTF). PCR positive samples were then sequenced prior to use for virus isolation. For quality control and lineage designation bioinformatic analysis was performed using the ARTIC workflow based on Nextclade and Pangolin v2023.06.10-1862208.

Details of all isolates used in this study, with their Spike mutations are detailed in Methods Table 1 below. Omicron sub-variants isolated at the Francis Crick Institute were collected from Legacy participants reporting acute symptomatic infection, following previously described active surveillance protocols^25^.

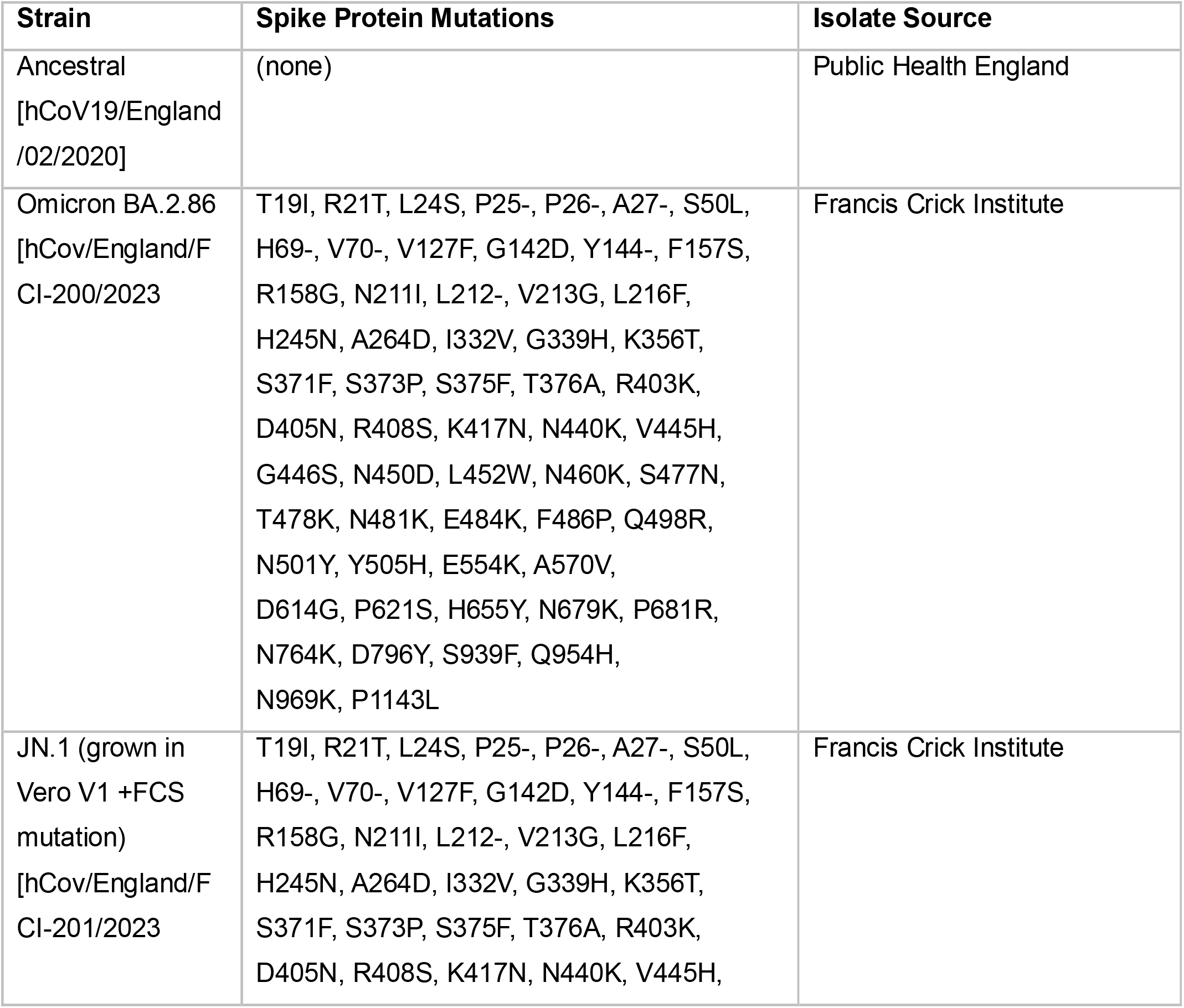
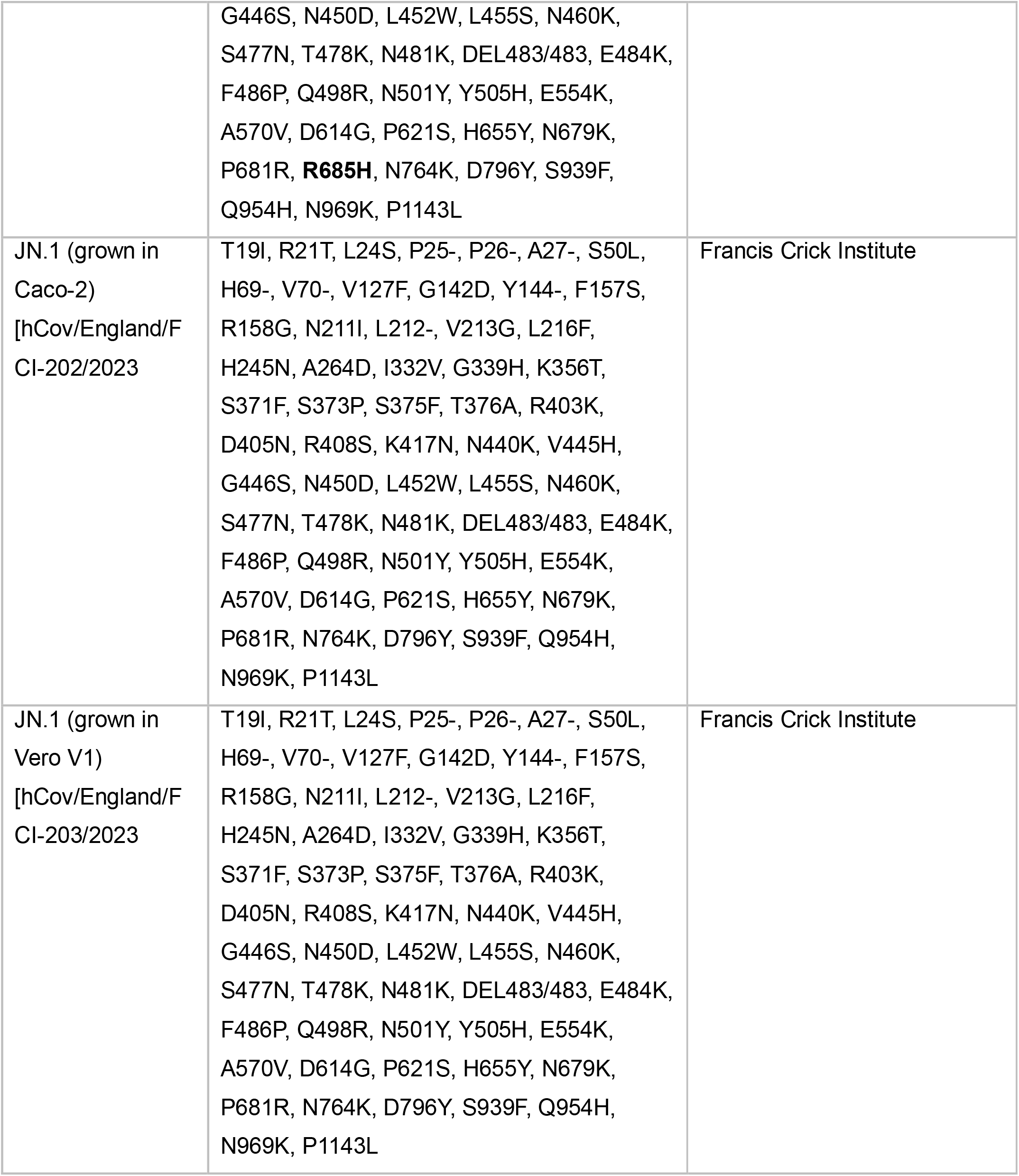

Methods Table 1. Summary of SARS-CoV-2 variants used, their Spike mutational profile and isolate source.

### High-throughput live-virus microneutralisation assay for serum samples

High-throughput live-virus microneutralisation assays for serum samples were performed as previously described^7,9,10,14,15,17,18^.

## Data analysis, statistics, and availability

Data used in this study were collected and managed using REDCap electronic data capture tools hosted at University College London^26,27^. Data were imported to R from REDCap prior to analysis as previously described^7-10,14,15^. Data were manipulated, analysed and visualised using *tidyverse* R packages^28^ including *dplyr* and *ggplot2*^29,30^. Summary descriptions of the cohort were generated using *gtsummary*^31^. Continuous data were reported as the median value and interquartile range (IQR) or the first and third quartiles (Q1; Q3). Statistical tests were conducted with the *rstatix* R package^32^.

Analysis of neutralising antibody titres in serum was performed as previously described without alterations using unpaired two-tailed Wilcoxon signed-rank tests^7-10,14,15,33^. Fold changes (FC) were estimated between groups with a 95% confidence interval (CI) with the *boot* R package using 5000 bootstrap resamples^34,35^.

Anonymised data (anonymised) and full R code to produce all figures and statistical analysis presented in this manuscript are freely available online on Github: https://github.com/DGreenwd/Crick-UCLH-Legacy-live-virus-microneutralization-for-SARS-CoV-2-variant-JN.1

## Ethics

The Legacy study was approved by London Camden and Kings Cross Health Research Authority (HRA) Research and Ethics committee (REC, reference 20/HRA/4717) IRAS number 286469 and 156 sponsored by University College London.

## Declaration of interests

CSw reports interests unrelated to this Correspondence: grants from BMS, Ono-Pharmaceuticals, Boehringer-Ingelheim, Roche-Ventana, Pfizer and Archer Dx, unrelated to this Correspondence; personal fees from Genentech, Sarah Canon Research Institute, Medicxi, Bicycle Therapeutics, GRAIL, Amgen, AstraZeneca, BMS, Illumina, GlaxoSmithKline, MSD, and Roche-Ventana, unrelated to this Correspondence; and stock options from Apogen Biotech, Epic Biosciences, GRAIL, and Achilles Therapeutics, unrelated to this Correspondence. DLVB reports discussions between the Crick and GSK for commercial antiviral testing, and grants to the Crick from AstraZeneca unrelated to this Correspondence. All other authors declare no competing interests.

## Consortium Authors

### Crick COVID serology pipeline

Ashley S Fowler, Murad Miah, Callie Smith, Mauro Miranda, Philip Bawumia, Harriet V Mears, Lorin Adams, Emine Hatipoglu, Nicola O’Reilly, Scott Warchal, Karen Ambrose, Amy Strange, Gavin Kelly, Svend Kjaer

### Legacy Investigators

Rupert CL. Beale, Padmasayee Papineni, Tumena Corrah, Richard Gilson

## Funding

This work was undertaken at UCLH/UCL who received a proportion of funding from the National Institute for Health Research (NIHR) University College London Hospitals Department of Health’s NIHR Biomedical Research Centre (BRC). EW, VL and BW are supported by the Centre’s funding scheme. This work further supported by the UK Research and Innovation and the UK Medical Research Council (MR/W005611/1, MR/Y004205/1, and MR/X006751/1 to EJC), and by the Francis Crick Institute which receives its core funding from Cancer Research UK (CC2166, CC1283, CC1114, CC2230, CC2060, CC2041, CC0102), the UK Medical Research Council (CC2166, CC1283, CC1114, CC2230, CC2060, CC2041, CC0102), and the Wellcome Trust (CC2166, CC1283, CC1114, CC2230, CC2060, CC2041, CC0102).

ECW, MW and NL are additionally supported by the PROVAC consortium via UK Research and Innovation (UKRI). DLVB is additionally supported by the Genotype-to-Phenotype National Virology Consortium (G2P-UK), Genotype-to-Phenotype 2 (G2P2-UK) and via UK Research and Innovation and the UK Medical Research Council. The funders of the study had no role in study design, data collection, data analysis, data interpretation, or writing of the report. The corresponding authors had full access to all the data and the final responsibility to submit for publication.

## Acknowledgements

The authors would like to thank all the study participants, the staff of the NIHR Clinical Research Facility at UCLH including Miguel Alvarez and Marivic Ricamara. We would like to thank the staff of the Scientific Technology Platforms (STPs) and COVID-19 testing pipeline at the Francis Crick Institute.

## Notes

https://github.com/DGreenwd/Crick-UCLH-Legacy-live-virus-microneutralization-for-SARS-CoV-2-variant-JN.1

