## Supplementary material for "Accurate evaluation of live-virus microneutralization for SARS-CoV-2 variant JN.1": FigureS1

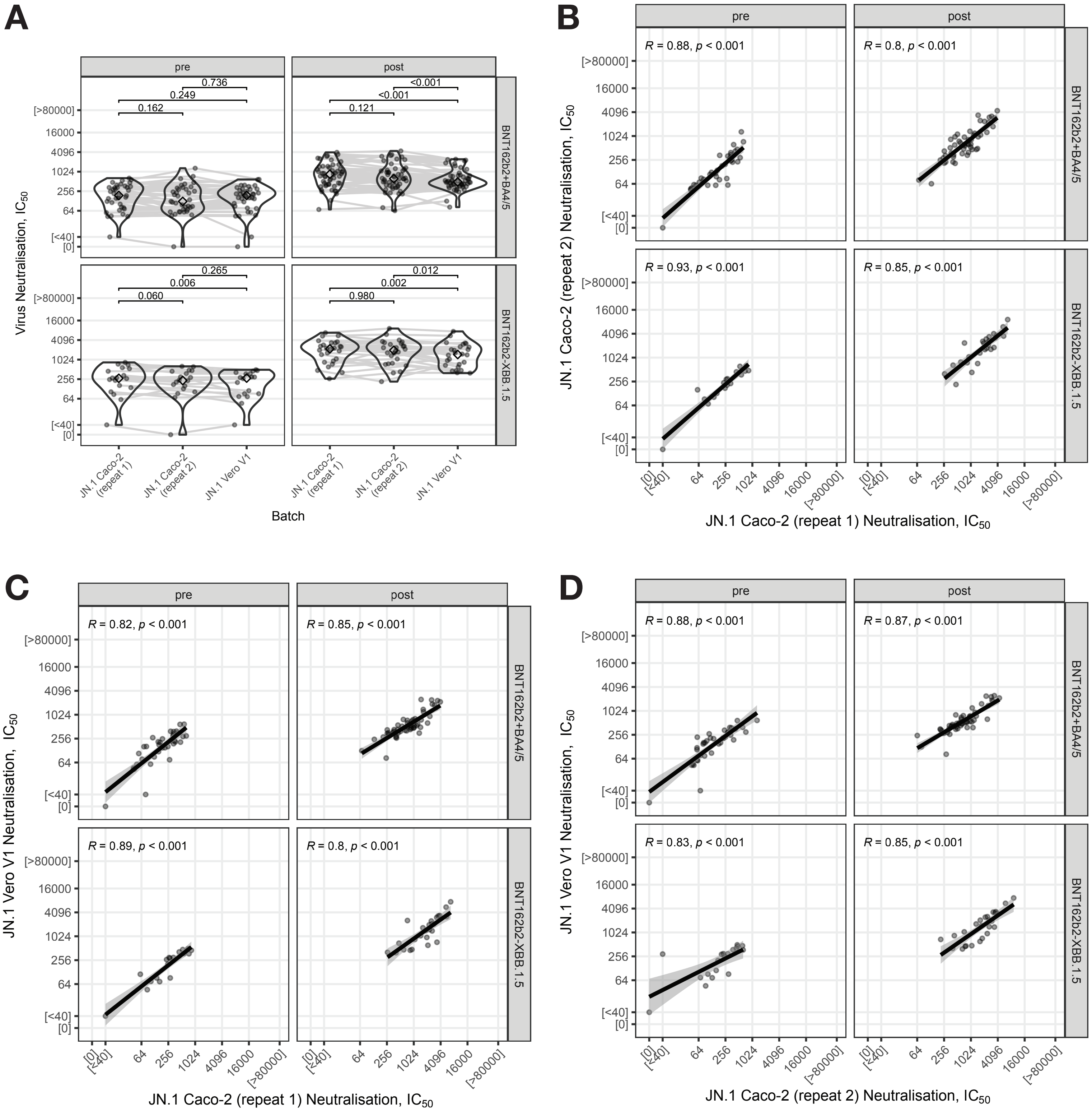


**Figure S1: Reproducibility and correlation between repeats of Caco-2 culture of JN.1 and Vero V1 culture of JN.1.** **(A)** Distribution of live-virus microneutralisation titres against SARS-CoV-2 JN.1 cultured in different cell lines are shown as the log2 of the IC50 for serum samples taken before or after a 5th dose of the bivalent BNT162b2 ancestral+BA.4/5 (top row) or BNT162b2-XBB.1.5 monovalent vaccine (bottom row). P values shown are from paired, two-tailed Wilcoxon tests, or McNemar's χ2 tests if the median of one group was more or less than the quantitative range of the assay (40–2560). FC = fold-change increase of nAbTs with 95% CIs in brackets. IC50 = 50% inhibitory concentration.
Correlation plots between live-virus nAbTs against SARS-CoV-2 JN.1 cultured in Caco-2 cells in two separate experiments **(B)**, and serum titres generated against JN.1 virus stocks in Vero V1 compared to the Caco-2 repeat 1 **(C)** and repeat 2 run within the same experiment **(D)**. P values and R coefficients of correlation are defined by the Spearman's comparison.
